# Automated Pipeline for Infants Continuous EEG (APICE): a flexible pipeline for developmental studies

**DOI:** 10.1101/2021.05.21.445085

**Authors:** Ana Fló, Giulia Gennari, Lucas Benjamin, Ghislaine Dehaene-Lambertz

**Author notes:** Declarations of interest: none. **Data statement:** The datasets analysed during the current study are available from the corresponding author.

## Abstract

Infant electroencephalography (EEG) presents several challenges compared with adult data. Recordings are typically short. Motion artifacts heavily contaminate the data. The EEG neural signal and the artifacts change throughout development. Traditional data preprocessing pipelines have been developed mainly for event-related potentials analyses, and they required manual steps, or use fixed thresholds for rejecting epochs. However, larger datasets make the use of manual steps infeasible, and new analytical approaches may have different preprocessing requirements. Here we propose an Automated Pipeline for Infants Continuous EEG (APICE). APICE is fully automated, flexible, and modular. Artifacts are detected using multiple algorithms and adaptive thresholds, making it suitable to different age groups and testing procedures without redefining parameters. Artifacts detection and correction of transient artifacts is performed on continuous data, allowing for better data recovery and providing flexibility (i.e., the same preprocessing is usable for different analyses). Here we describe APICE and validate it using two infant datasets of different ages tested in different experimental paradigms. We also tested the combination of APICE with common data cleaning methods such as Independent Component Analysis and Denoising Source Separation. APICE uses EEGLAB and compatible custom functions. It is freely available at https://github.com/neurokidslab/eeg_preprocessing, together with example scripts.

## 1. INTRODUCTION

Electroencephalography (EEG) provides a non-invasive, direct, and low-cost measure of neural activity with a high temporal resolution, making EEG a valuable tool for developmental cognitive studies. Two problems are encountered with this technic. First, the EEG signal is unavoidably contaminated by many artifacts from different sources, such as environmental factors (e.g., line noise), physiological phenomena (e.g., ocular artifacts, heartbeats, muscle activity), and motion artifacts, whose amplitude is often much larger than the neural signal. Second, the neural signal of specific cognitive processes is lost among many other computations occurring at the same time, which furthermore spatially overlaps due to the wide diffusion of the electrical fields. One successful solution to isolate a cognitive process is to average across many trials to recover a reproducible neural activity time-locked to the stimulus presentation, i.e., the event-related potential (ERP). For the averaging method to be successful, many trials are needed, and those trials should not be contaminated by high amplitude changes, whose impact on the average could not be eliminated without thousands of trials. Other EEG analysis techniques (e.g., decoding, time-frequency analyses) have similar constraints on a high number of trials, without large amplitude changes. However, these requirements are at odds with the usual testing circumstances in infants, which are short and often heavily contaminated by motion, thus calling for a different approach to obtain a sufficiently good signal despite these challenging conditions. While simple steps as filtering can remove some artifacts as line noise, the correction of physiological artifacts requires more sophisticated methods (Islam et al., 2016). Fortunately, the high redundancy of the signal in time and space due to the diffusion of the electric field enables the implementation of different signal reconstruction techniques (Jiang et al., 2019).

Several pipelines and artifact removal algorithms have been developed for adult EEG (e.g., PREP (Bigdely-Shamlo et al., 2015), Automagic (Pedroni et al., 2019), FASTER (Nolan et al., 2010), ADJUST (Mognon et al., 2011), MARA (Winkler et al., 2011)). However, they are not adapted to infants challenging data. The correction methods currently available for physiological artifacts require long recordings without high amplitude artifacts (Onton & Makeig, 2006); thus, they are not directly applicable to infant data. Second, the power spectrum of the EEG signal (Eisermann et al., 2013; Marshall et al., 2002) and the properties of the evoked responses (Kushnerenko et al., 2002; Nelson & Monk, 2001) evolve throughout development due to maturational changes. More specifically, the background activity is rich and ample in low frequencies, and the signal variability between trials is higher in infants than in adults (Naik et al., 2021). Third, exogenous artifacts vary according to infants’ age (e.g., fewer blinks and less motion in younger infants). Lastly, most infant datasets do not include electrocardiogram (ECG) and electromyogram (EMG) recordings used in adults’ pipelines to identify non-neural artifacts.

Because of all the factors mentioned in the previous paragraph, the methods developed for adult EEG are ill-suited for infant studies, and no agreement has been reached on the most appropriate preprocessing procedures for infant EEG. Traditional preprocessing usually relies on the manual identification of non-functional channels and data segments contaminated by motion artifacts, which are subsequently discarded. However, high-density recording systems (64, 128, 256 electrodes) and longer recording sessions make this approach time-consuming and inefficient, revealing the need for automated pipelines. These are usually based on fixed thresholds below which the voltage must remain. Non-functional channels are either rejected or interpolated, but no additional correction of non-neural artifacts is applied (Adibpour et al., 2018; Friedrich et al., 2015; Kabdebon & Dehaene-Lambertz, 2019). Although straightforward, this method requires thresholds to be set arbitrarily, weighing the risk of a high rejection rate that might leave too few trials to obtain an ERP free of background activity and a low rejection rate that might retain an artifactual signal in the ERP. Because signal’s amplitude depends on the distance between the channel and the reference, fixed thresholds are more or less able to detect artifacts, with the risk of being too sensitive for distant electrodes and not enough for close electrodes. Furthermore, given the changes in EEG amplitude as a function of age and sleep-wakefulness stages, fixed thresholds need to be re-adjusted for each particular dataset. Finally, the detection of artifacts is done on segmented data, usually short epochs, limiting its applicability on methods requiring longer recordings.

More complex paradigms are implemented nowadays (Friedrich et al., 2015; Kabdebon & Dehaene-Lambertz, 2019) to explore infants’ rich cognition. The impatience of young subjects, the lack of verbal instructions, and the common need of having familiarization periods make the amount of data in pertinent conditions scarce, making maximum data retention without quality lost crucial. Furthermore, new analysis techniques with different requirements are now being implemented. For example, frequency tagging may require segmenting the data in longer epochs (e.g., de Heering & Rossion, 2015; Kabdebon et al., 2015). Researchers may also want to retain as much data as possible for multivariate decoding (Gennari et al., 2021).

Two demands that can be hard to reconcile with a high degree of artifact contamination. Moreover, different types of analyses might be used on the same dataset, requiring the same data to be processed several times. Two papers proposing pipelines for developmental data have recently been published, HAPPE (Gabard-Durnam et al., 2018) and MADE (Debnath et al., 2020). Both pipelines focus on the implementation of ICA for the removal of non-neural artifacts. While they provide different strategies for a proper application of ICA on infants’ data, they do not provide a specific way to deal with the high amount of motion artifacts present in infants’ recordings, which can drastically affect ICA or any other preprocessing step.

Here we propose an Automated Pipeline for Infants Continuous EEG (APICE), in which automatized artifact detection is performed on the continuous data before any further preprocessing step. APICE provides high versatility and flexibility and offers good data recovery, and ensures data quality. We have developed APICE to have a standard preprocessing procedure, flexible enough to (1) be appropriated for a broad range of analyses; (2) applicable on different developmental populations; and (3) able to recover as much good quality data as possible. Relatively to previous work, the key innovations we propose with APICE are: (1) an iterative artifact detection procedure based on multiple algorithms applied on continuous data, (2) the use of automatically adapted thresholds, and (3) the correction of transient artifacts on continuous data. The use of multiple algorithms and adaptive thresholds makes APICE applicable without modifications across different ages and protocols. It also makes it easily adaptable to adult EEG datasets. The early detection of artifacts in the continuous data enables the experimenter to decide how to deal with artifacts before applying further preprocessing steps, preventing leaks on non-contaminated parts of the data. Additionally, the detection and correction of artifacts on continuous data increase data recovery by avoiding rejecting data segments containing transient voltage jumps. Finally, APICE is highly flexible, allowing using the same preprocessed data for many kinds of analysis. Note that we build this pipeline to suit our own needs, that is to say, to use high-density nets to explore infants’ cognition. Processing clinical data with a few electrodes has neither the same goal of high robustness at the individual level nor the same spatial redundancy to interpolate data. Nevertheless, we believe that most of the solutions we propose in APICE are usable in many experimental and clinical situations keeping in mind the signal features and the purpose of the EEG recording. It is why all parameters are adjustable even if we provide default values based on our experience.

In this paper, we first describe the general logic behind the pipeline and its different steps. We then provide validation of APICE by comparing it with a standard preprocessing pipeline on two datasets with very different properties. We also compare APICE with a reduced version of it. In the latter version, we kept the automatic detection of artifacts in continuous data but removed the correction of transient artifacts to evaluate the effect of both processes in terms of data quality and data recovery. Finally, we evaluate whether incorporating additional data cleaning methods described in the literature provides any improvement. Specifically, we tested ICA and Denoise Source Separation (DSS) (De Cheveigné & Parra, 2014; de Cheveigné & Simon, 2008).

APICE is implemented in MATLAB, and it uses the EEGLAB toolbox and custom functions compatible with the EEGLAB structure. It is freely available at https://github.com/neurokidslab/eeg_preprocessing, together with example scripts. APICE is modular, allowing functions to be easily recombined to meet different requirements. APICE also provides toolkit functions to modify events, correct event timings using Digital Input events (DINs), add information about trial features, and obtain average ERPs.

## 2. PIPELINE GENERAL DESCRIPTION

APICE uses EEGLAB (Delorme & Makeig, 2004) functions for standard processing steps (e.g., importing the data, filtering, epoching) and includes new functions for more specific steps. The crucial additions in APICE are (1) the identification of motion artifacts on continuous data using relative thresholds; (2) the correction of artifacts in the continuous data when they involve a few channels or affect a brief period; (3) the possibility to define contaminated samples and non-functional channels based on the rejected data; (4) the definition of bad epochs based on the amount of rejected data within the epoch in term of time samples and electrodes.

Additionally, APICE includes functions which allow to apply other standard data cleaning methods. We provide a function to perform ICA, which omits the samples and channels previously identified as containing artifacts, uses Wavelet-thresholding before performing the ICA, and automatically identifies components associated with artifacts using iMARA (Marriot Haresign et al., 2021). We also provide a function to apply DSS, a method to clean ERP proposed by de Cheveigné and colleagues (De Cheveigné & Parra, 2014; de Cheveigné & Simon, 2008).

All the functions and steps are modular. While we propose the different steps to be performed in a specific order to obtain optimal results, the functions can be re-combined according to particular needs. From our experience in infants EEG data, we propose the following steps for preprocessing infant EEG data (Figure 1).

1. Importing the data to EEGLAB
2. Minimal filtering of the data
3. Detecting artifacts
4. Correcting artifacts when possible
5. Re-detecting artifacts
6. Applying ICA (optional)
7. Epoching
8. Rejecting bad epochs
9. Applying DSS (optional)
10. Reference averaging, data normalization, baseline correction (optional)

**Figure 1.**
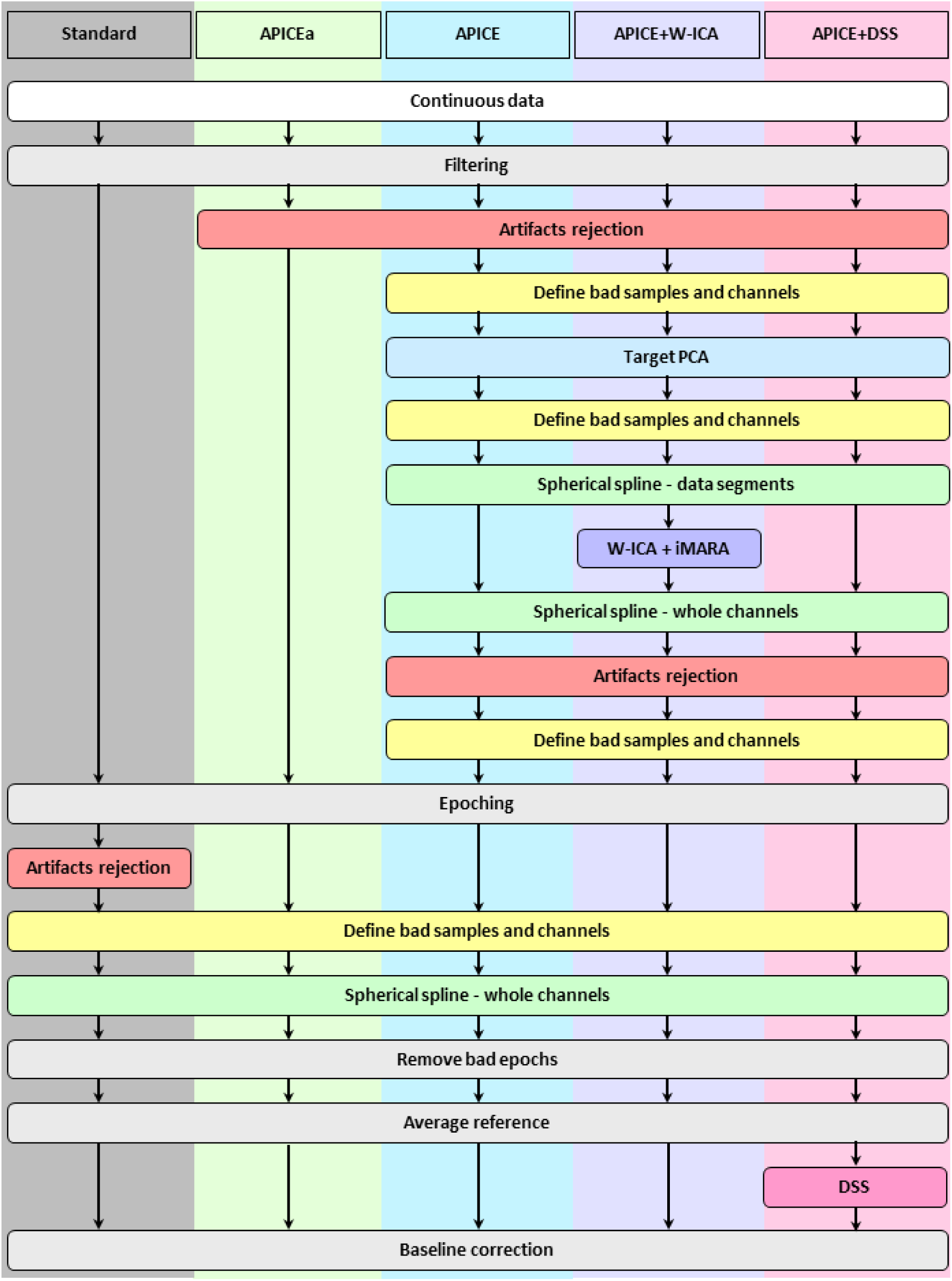
Schematic description of different preprocessing pipelines. Standard corresponds to a common basic preprocessing pipeline usually used on infant experiments. APICEa is a reduced version of our pipeline in which artifacts detection is performed on continuous data using multiple algorithms and relative thresholds (see section 2.2). APICE is our full preprocessing pipeline with the correction of transient artifacts on continuous data (see section 2.4). APICE +W-ICA is APICE with the addition of ICA. APICE+DSS is APICE with the addition of DSS.

Note that the steps after epoching (step 7) are specific to ERP analyses or other analyses involving evoked activity (i.e. an averaging process across trials to recover reproducible time-locked activity).

### 2.1. Filtering

As the first step of the preprocessing, data are filtered to remove common environmental artifacts. We use a low-pass filter below the line noise frequency (e.g., with a line noise at 50 Hz, we low-pass filter the data at 40 Hz). Unless high-frequency activity needs to be investigated, low-passing filtering is, in our experience, the most effective way to remove line noise.

High pass filtering enables the removal of drifts and slow activity in the data. Low-frequency noise strongly contaminates high impedance EEG recordings, mainly due to skin potential (Kappenman & Luck, 2010). Furthermore, slow waves are common in very young infants’ EEG recordings (Eisermann et al., 2013; Marshall et al., 2002; Selton et al., 2000), due to immaturity but also partially due to the brain’s movement in the skull following respiration in young infants. For most EEG analysis, it is imperative to reduce this contamination. However, extensive filtering (above 0.1 Hz) can introduce critical distortions in the data (de Cheveigné & Nelken, 2019). While alternatives to high-pass filtering exist, such as local detrending methods (de Cheveigné & Arzounian, 2018), we have not observed improved results. Therefore, at this early stage of preprocessing, we apply a high pass filter at 0.1 Hz, which should not introduce critical distortions in the data but remove the main drifts.

We use the *pop_eegfiltnew* EEGLAB function, which uses the FIRfilt plugin, to perform a non-causal Finite Impulse Response (FIR) filter. We first apply a low-pass filter at 40 Hz with a transition band of 10 Hz. We then apply a high-pass filter at 0.1 Hz with a transition band of 0.1 Hz.

It is worth noticing that high-pass filters should only be applied on non-epoch data to avoid edge effects. While low-pass filters can be applied at any stage of the preprocessing, further high-pass filter should always be performed on the continuous data.

### 2.2. Artifacts detection

One of the main differences of APICE with other pipelines is the detection of artifacts. Artifacts are identified on the continuous data before average referencing, and we use adaptive rather than absolute predefined thresholds, allowing the parameter to be automatically defined per subject and electrode. As we discuss below, these features are decisive for an adequate and versatile preprocessing pipeline for infant recordings.

Non-working electrodes have a response radically different from the rest of the channels, with amplitudes much higher or lower than expected. However, in unipolar recordings (i.e., the signal is recorded as the difference of potential between each electrode and a single reference), the amplitude of the signal varies in function of the distance of each electrode to the reference electrode—with electrodes closer to the reference having smaller amplitudes than more distant ones. Thus, absolute thresholds classically used in standard pipelines differently penalize each channel. If the signal’s amplitude is already large, a slight supplementary increase might be interpreted as an artifact, whereas the same deviation in electrodes close to the reference might remain undetected. Therefore, the same noise level is not similarly detected across electrodes, and the percentage of recording rejected in each channel might be very different at the end of the process.

A solution to homogenize the voltage in high-density recordings is to measure the potential relative to an ideal reference, such as the average reference (Bertrand et al., 1985). The integrated scalp potential must be null and can be approximated by the average over numerous homogeneously distributed channels. Average referencing achieves a null integrated potential by subtracting the average voltage at each time point to each electrode. Nevertheless, to avoid affecting functional channels with contaminated data through the average process, the data must be clean.

We thus face a circular problem: to estimate non-functional channels based on their amplitude, we need to average referenced the data, but to obtain a proper estimation of the average reference, we need clean data. To overcome this problem, we identify artifacts and non-working electrodes before estimating the average reference. We do so by using multiple algorithms sensitive to different signal features. Some algorithms are independent of the signal amplitude, while in others, thresholds are adapted to the signal level recorded by each electrode. Thus, all are applicable before average referencing.

Another advantage of adaptive thresholds is that they do not have to be customized to each population or testing procedure (i.e. age group, resting-state or active task). The amplitude and properties of the signal radically change during development, and between-subjects variability is considerable. Therefore, using adaptive thresholds is an essential feature for infant EEG.

The detection of artifacts in continuous data rather than in segmented epochs also has its own advantages. First, the properties of the EEG recording and the threshold for the different artifacts can be better estimated. Second, some transient artifacts that may lead to the rejection of the epoch can be corrected, facilitating the recovery of more data. Third, it provides more flexibility, enabling the implementation of multiple types of analysis using a common preprocessing pipeline.

All algorithms compute a measure for each sample or in a sliding time window. Then, the algorithms reject the data when the measure is above or/and below a threshold. In all cases, it is possible to use absolute thresholds but our advice is to use relative thresholds. Relative thresholds are determined based on the distribution of the measure throughout the entire recording. More precisely, a threshold is computed as, *Thresh* = *Q*_3_ + *k* × (*Q*_3_ – *Q*_1_), and/or *Thresh* = *Q*_1_ − *k* × (*Q*_3_ – *Q*_1_), where *Q*_1_ and *Q*_3_ are the first and third quartiles of the distribution, and *k* should be provided as an input (by default 3). Because the measures used by the algorithms usually have a normal distribution, this threshold definition successfully identifies extreme values. A single threshold can be defined for all electrodes, or individual thresholds can be established per electrode. We also provide the option of computing the algorithms in data z-scored per channel or average reference. However, the output data is never modified.

The algorithms define for each sample and channel if the data is good or contains artifacts, and the information is stored in a logical matrix of the size of the recording (i.e., channels x samples x epochs (equal to 1 in case of continuous recording)), where a true value indicates the presence of artifacts.

Different algorithms are sensitive to different artifacts in the data. Using a collection of algorithms and methods enables a large and complete detection of artifacts of all kind in the data. Some of the algorithms are particularly sensitive to the detection of non-functional channels. Specifically, one algorithm looks at the power spectrum of the different channels across frequency bands, and another detects channels with very low activity correlation with other channels. Other algorithms are sensitive to motion artifacts. They detect when the signal’s amplitude, temporal variance, or running average are too high. Another algorithm identifies when the signal changes too rapidly and serves to detect jumps or discontinuities in the signal. A final group of algorithms rejects, or re-includes data, according to the rejection matrix and its “salt and pepper” structure. For example, they reject segments of data if they are too short and between rejected segments, or re-include very short rejected data segments that were previously rejected. A full description of all the algorithms and functions is provided in Appendix A.

In APICE, we detect artifacts through multiple cycles of rejection and the data rejected in one cycle is no longer entered into the signal estimation used to construct adaptive rejection thresholds for subsequent cycles. These multiple cycles allow a progressive skimming of the signal. After each cycle, we re-include rejected segments shorter than 20 ms, and we reject good data segments between bad segments shorter than 2 s. We propose the following rejection cycles:

- Rejection cycle 1: we run the algorithms that primarily identify channels not working based on their power spectrum and their absence of correlation with other channels.
- Rejection cycle 2: we reject all data with an amplitude higher than 500 mV (non-average referenced data). This absolute threshold is very high and is only used to avoid taking into account very large amplitude data. This step accelerates the reiteration procedure but can be skipped without substantial changes.
- Rejection cycles 3a and 3b: we apply twice the algorithms sensitive to motion artifacts using relative thresholds per electrode and non-average referenced data. Specifically, the algorithms reject data with a too high amplitude, too high variance, or too high running average relative to themselves.
- Rejection cycles 4a and 4b: we apply the same algorithms as in the third cycle twice, but this time on average referenced data and using one relative threshold across all electrodes.
- Rejection cycles 5a and 5b: we detect fast transient changes in the signal, once using one threshold per electrode on non-average reference data and one on average reference data and defining a single threshold across all electrodes.

After artifact detection, we correct artifacts (see section 2.4). Then, the rejection matrix is reset, and the artifact detection algorithms are re-run to detect artifacts in the “clean” data. Because non-functional channels and jumps in the signal were already corrected when possible, we only re-run algorithms to detect motion artifacts (rejection cycles 2 to 4).

### 2.3. Definition of bad samples and channels

The rejection matrix is a logical matrix of the size of the data indicating outlier points. However, even if some data points are not identified as outliers, they can still contain artifacts. It is the case of short “good” periods between “bad” periods and when most of the channels, but not all, are considered “bad”. Thus, we also defined bad-times (BT) to refer to time samples strongly contaminated by artifacts (most of the channels were identified as “bad”) and bad channels (BC) to refer to non-functional channels (most of the times identified as “bad”). These definitions allow to easily exclude from future analysis samples containing large artifacts that cannot be corrected and reconstruct non-functional channels. The functions used to define BT and BC are described in Appendix B. In Figure 2, we present an example of a rejection matrix and the definition of BT and BC.

**Figure 2.**
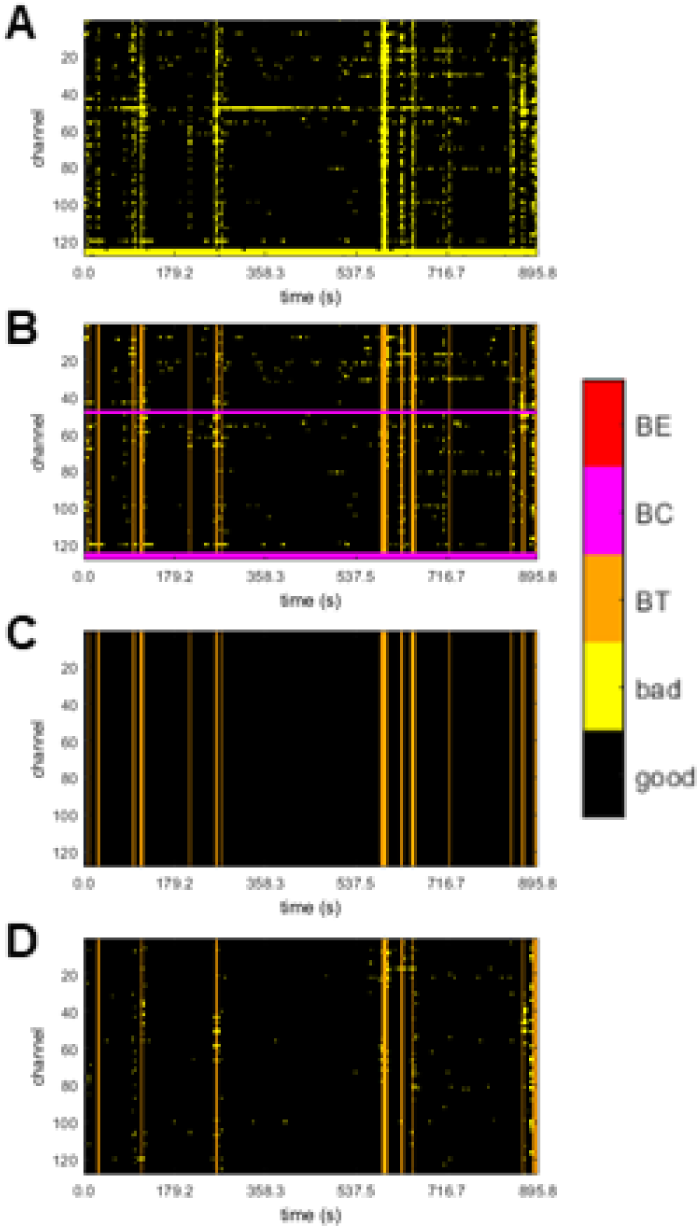
Example of the rejection matrixes (channels x time-samples) for one subject obtained during successive stages of the preprocessing. BE = Bad Epochs, BC = Bad Channels, BT = Bad Times. **(A)** Rejection matrix after the artifacts detection. **(B)** Identification of bad channels (pink) and samples (orange). **(C)** Rejection matrix with bad times and channels after the interpolation of localized artifacts (bad channels and transient artifacts). Notice that all rejected samples, except bad times, have been corrected. **(D)** Rejection matrix after applying again the artifacts detection algorithm. Most of the corrected artifacts are no longer identified as artifacts.

In our practice, we define periods in which artifacts affect more than 30 % of functional electrodes as “bad times”. We consider as good, bad segments shorter than 100 ms. We mark as bad the 500 ms before and after any bad time segment to avoid correcting non-functional channels close to motion artifacts and considering data samples surrounding motion artifacts, which tend to be partially contaminated. We reject good-time segments shorter than 1 s.

The channels presenting artifacts during more than a certain proportion of the good times are rejected. Bad channels can be defined during each epoch (BC) or during the whole recording (BCall). This distinction is only important when working with epoched data. In continuous data, we define bad channels as those that are not functional for more than 30% of the good times. We defined bad channels in a given epoch as those showing an artifact lasting more than 100 ms during good times.

### 2.4. Correction of localized artifacts

Artifact correction is a critical point because it implies data reconstruction. Data segments containing large artifacts (i.e., motion artifacts affecting most electrodes during a period) are very common in infant recordings but unfortunately cannot be reconstructed and need to be discarded. Thus, the best we can aim is to identify them accurately. These periods correspond to the bad times defined in the previous section. On the contrary, other types of artifacts can be corrected. We have already described the removal of electrical noise by filtering. Additionally, many methods exist to correct physiological non-neural sources (Islam et al., 2016; Jiang et al., 2019). We will specifically discuss the use of ICA to clean this type of artifact in the next section. This section discusses data reconstruction when artifacts are localized in either time or space (channels). A detailed description of the function for data correction is presented in Appendix C.

Transient artifacts, like jumps and discontinuities, frequently contaminate the EEG signal. To remove this type of artifact, we apply a target PCA and remove the first components. The underlying assumption is that the majority of the variance can be attributed to the artifact during the artifact period. This approach has already been implemented on Near-Infrared Spectroscopy data (Yucel et al., 2014). Contrary to blind source separations methods, like ICA or PCA applied to the whole recording, the PCA is here restricted to segments with artifacts. Thus, it limits the undesired removal of neural activity. We used it for segments shorter than 100 ms, and we removed the first components carrying 90 % of the variance. The time limit for the length of the artifacts circumscribes the correction to jumps in the signal and other fast non-neural sources such as heartbeat (see Figure 3).

**Figure 3.**
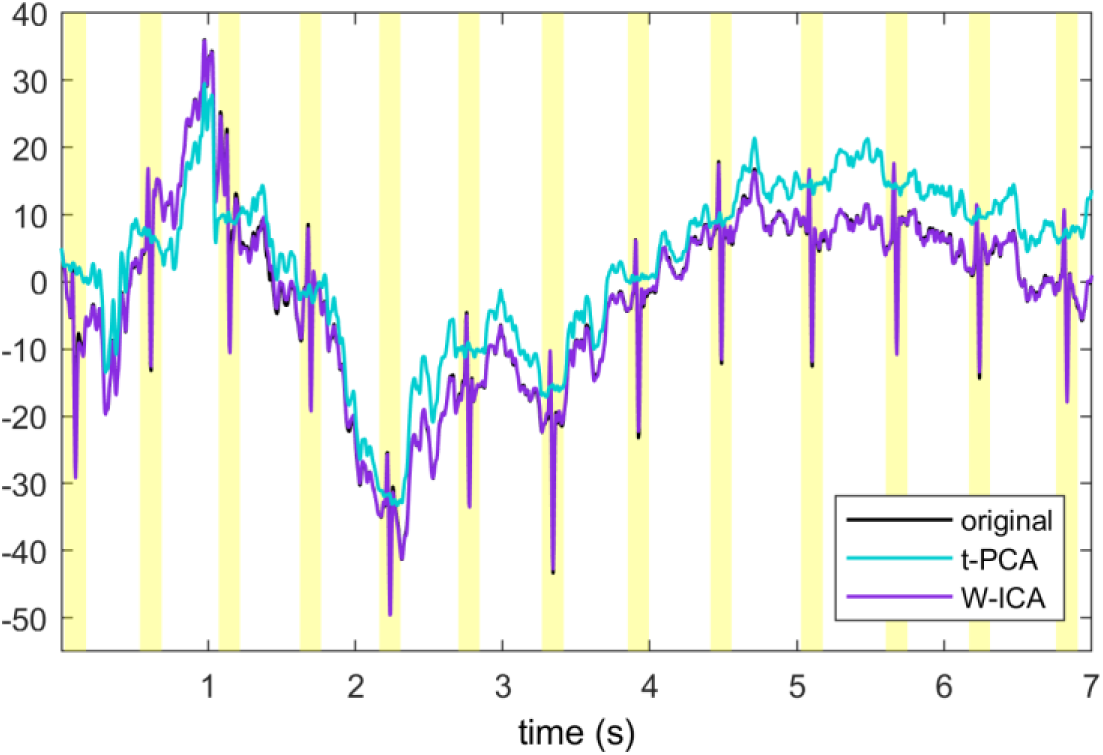
Example of heartbeat artifact correction by target-PCA and ICA. The shaded area represents the samples identified as having artifacts, and on which target-PCA is applied.

Another possible scenario is that artifacts affect a small number of electrodes as in the case of non-functional electrodes. A widely used reconstruction method for these cases is the spatial interpolation of the channels using a spherical spline (Perrin et al., 1989). In adult experiments, electrodes are identified as non-working during the whole recording and eventually spatially interpolated. With infant high-density systems, it is common that some channels stop working during limited periods (e.g., the infant touches some electrodes, making them lose contact, or after a child’s movement, the electrode moves before returning to its original position). To account for these scenarios and recover as much data as possible, we reconstruct the channels identified as bad during the whole recording and during restricted periods, by spatial interpolation. It is worth noticing that while the reconstruction of non-functional channels is a common practice that can simplify the between-subjects analysis, the signal is estimated as a weighted sum of the good-electrodes. Thus, it does not add any new information, and the dataset becomes rank deficient.

In APICE we apply the artifacts correction as follows. First, we correct transient artifacts using target PCA. Second, we spatially interpolate channels not working during a period. We restrict this interpolation to good times, that is to samples where no more than 30% of the channels are excluded. The underlying assumption is that bad samples contain too strong artifacts affecting too many channels; hence, the spatial interpolation is not effective. Finally, we interpolate channels indicated as non-functional during the whole recording. By applying the functions in this order, transient artifacts are corrected before the non-functional channels, minimizing the use of contaminated data to reconstruct non-functional channels. Figure 2C shows the rejection matrix after interpolation. Observe how bad data remains only during bad times.

### 2.5. Independent Component Analysis

ICA is commonly applied on multichannel recordings of EEG data to remove physiological non-neural components (e.g., eye blinks, eye-motions, muscle activity, heartbeat). The recording is decomposed in temporally independent components (IC). Then, the components related to artifacts are identified either automatically or manually by their topography, temporal profile, and power spectrum and removed from the data. Ideally, each IC is related to a neural or non-neural source of the recorded activity. Therefore, ICA allows removing specific artifacts (sources) from the data without discarding the EEG segments affected by that artifact. It is worth noticing that removing a specific IC alters the entire recording, with the risk of eliminating genuine neural activity. The successful application of ICA for EEG data cleaning depends on (1) an appropriate separation of non-neural sources in distinct components and (2) their proper identification.

ICA is a common practice in the preprocessing of adult data. However, its implementation in infant data is not widespread, and its benefits are not clear. The poor performances of the method are partially due to the nature of infant EEG data. Young infants’ recordings contain more slow waves than adult data (Eisermann et al., 2013; Marshall et al., 2002), ERPs are less precise in time (Kushnerenko et al. 2002) and more variable (Naik et al., 2021). The nature of the artifacts is also different in infants and adults. Adults are often quiet and attentive, with a low voltage EEG. Thus, artifacts are rare and have a clearly different signal, contrary to infants. All these factors degrade a successful separation of neural and non-neural sources. A second reason for these lower performances is that algorithms for the automatic identification of artifact-related IC have been mainly developed for adult data. Recently, some algorithms have been adapted for infant data. For example, the ADJUST algorithm (Mognon et al., 2011) has been adapted into the adjusted-ADJUTS (Leach et al., 2020), and the MARA algorithm (Winkler et al., 2011) into iMARA (Marriot Haresign et al., 2021). While these modified algorithms work better on infant data, they rely on optimizations done on adult training sets. Additionally, the continuous developmental change of the signal (e.g., changes in the power spectrum profile and ERP due to maturation of the neural circuits, changes in the diffusive properties of the skull due to its maturation) may hinder the classification and degrade the algorithm’s performance.

A good ICA decomposition requires several considerations. Previous work shows that to obtain a reliable separation in IC, the data has to be high-pass filtered at least at 1 Hz and should not contain high amplitude noise (e.g., motion artifacts) (Winkler et al., 2015). However, high-pass filtering the data may not be suitable for many EEG analyses (e.g., ERPs are distorted, and slow waves may be lost). We implemented a standard solution consisting of high-pass filtering and applying ICA to a copy of the data. In this way, the non-neural components are estimated and subtracted from the original data (Debnath et al., 2020).

To avoid having high amplitude noise on the data, we restricted the ICA to the samples identified as good times and set at zero the remaining data points marked as artifacts. Additionally, to remove potentially remaining transient high amplitude artifacts, we performed a wavelet-thresholding on a first ICA (Geetha & Geethalakshmi, 2011; Johnstone & Silverman, 1997). On the clean data, we applied ICA again. The use of wavelet-thresholding on a first ICA decomposition improves the final ICA decomposition (Gabard-Durnam et al., 2018; Rong-Yi & Zhong, 2005).

Another important consideration is that the recording should fulfill *m* ≥ 30 × *n*^2^, where *m* is the number of samples and *n* the number of channels (Onton & Makeig, 2006). Infants’ recordings are generally not long enough to guarantee this condition. There are two alternatives to overcome this issue. One is to reduce the analysis to a subset of channels. The second possibility is applying PCA and retaining only the first components to reduce the problem’s dimensionality.

We do not regularly apply ICA in APICE (see the pipeline validation section for further discussion). When we apply it, data are high-pass filtered at 2 Hz, and we use PCA first to reduce the dimensionality of the problem. We have noticed that with high-density nets, the variance lost by keeping only the first ∼50 components is minimal, and results are more accurate than by reducing the number of channels. With a smaller number of channels, the spatial resolution decreases, and we have observed a considerably worst performance of the classification algorithms. Moreover, reducing the number of channels implies either not analyzing some of them or analyzing the data in multiple loops. Therefore, we opted to use PCA instead for short recordings.

Notice that for ICA, the signals need to be independent. Thus, in principle, ICA should be applied before the spatial interpolation of the non-functional channels. However, if PCA is applied first to reduce the dimensionality, ICA can be performed before or after interpolating non-functional channels. We use the iMARA algorithm (Marriot Haresign et al., 2021) for the automatic identification of components associated with non-neural artifacts. In Appendix D we describe the function that performs ICA in APICE.

### 2.6. Definition of bad epochs

Once continuous recordings are segmented into epochs, we defined bad epochs that should not be considered in the following analyses. An epoch is defined as bad if any of the following three criteria is present: 1) it contains any bad time; 2) it contains more than 30 % of bad channels; 3) if more than 50 % of the data was interpolated. Note that 30 % is the limit in the proportion of channels to define bad times; thus, the first two criteria overlap. The function to define bad epochs is described in Appendix E.

### 2.7. Denoising Source Separation (DSS)

Spatial filters are linear combinations of the sensors designed to partition the signal between components carrying the signal of interest from non-interest. In the particular case of the DSS, the spatial filter is designed to select components carrying evoked activity, meaning activity that is reproducible across trials, from non-evoked activity. Thus, the method has been proposed as an alternative to clean ERPs (De Cheveigné & Parra, 2014; de Cheveigné & Simon, 2008). It is worth noticing that this data cleaning method is specific to the study of evoked activity because the activity that is not phase-locked to the stimuli is partially removed.

In Appendix D, we describe the function that we provide in APICE to perform DSS.

### 2.8. Preprocessing report

Within the EEGLAB structure, we register a report containing the size of the data, the amount of rejected data, and the amount of interpolated data after each data processing step. More specifically, we retain the number of channels, samples, epochs, the number of rejected and interpolated points, and the number of bad times, bad channels, and bad epochs. We also provide a function that prints a table in the command window and a text file to summarize these measures: The information is collected for all subjects at critical points during the pipeline. Specifically, it summarizes the rejection percentages before data epoching, before the rejection of bad epochs, and at the final stage. This summary information should help the experimenter to evaluate the pipeline’s performance and to detect possible problems.

A description of these functions can be found in Appendix E. As an example, the report for the analysis performed using the APICE pipeline on dataset 1 is shown in Table 1.

**Table 1.** Example of a preprocessing report. The summary table corresponds to Dataset 1 after the rejection of bad epochs. For each subject it shows the total number of channels (Ch), the total number of samples per epoch (Smpl), the total epochs (Ep), the percentage of bad data (BCT), the percentage of interpolated data (CCT), the percentage of bad times (BT), and the percentage of bad channels (BC). Notice that an analog table is also generated before rejecting the bad epochs.

| Subject | Ch | Smpl | Ep | BCT(%) | CCT(%) | BT(%) | BC(%) |
| --- | --- | --- | --- | --- | --- | --- | --- |
| Session 20180926 S02.set | 128 | 1000 | 214 | 0.001 | 8.2 | 0 | 0.00 |
| Session 20180927 S03.set | 128 | 1000 | 193 | 0.005 | 14.2 | 0 | 0.00 |
| Session 20180927 S04.set | 128 | 1000 | 198 | 0.004 | 11.5 | 0 | 0.00 |
| Session 20180927 S05.set | 128 | 1000 | 189 | 0.003 | 17.7 | 0 | 0.00 |
| Session 20181003 S06.set | 128 | 1000 | 182 | 0.002 | 7.3 | 0 | 0.00 |
| Session 20181003 S07.set | 128 | 1000 | 205 | 0.002 | 9.5 | 0 | 0.00 |
| Session 20181004 S08.set | 128 | 1000 | 212 | 0.001 | 8.8 | 0 | 0.01 |
| Session 20181004 S09.set | 128 | 1000 | 190 | 0.002 | 15.2 | 0 | 0.00 |
| Session 20181004 S10.set | 128 | 1000 | 213 | 0.004 | 8.5 | 0 | 0.05 |
| Session 20181005 S11.set | 128 | 1000 | 208 | 0.002 | 8.2 | 0 | 0.00 |
| Session 20181005 S13.set | 128 | 1000 | 202 | 0.001 | 13.5 | 0 | 0.00 |
| Session 20181022 S14.set | 128 | 1000 | 201 | 0.001 | 8.2 | 0 | 0.00 |
| Session 20181022 S15.set | 128 | 1000 | 198 | 0.003 | 16.0 | 0 | 0.05 |
| Session 20181022 S16.set | 128 | 1000 | 175 | 0.003 | 19.2 | 0 | 0.00 |
| Session 20181023 S18.set | 128 | 1000 | 208 | 0.001 | 12.6 | 0 | 0.00 |
| Session 20181023 S20.set | 128 | 1000 | 171 | 0.001 | 11.0 | 0 | 0.00 |
| Session 20181024 S21.set | 128 | 1000 | 188 | 0.004 | 12.3 | 0 | 0.04 |
| Session 20181024 S22.set | 128 | 1000 | 188 | 0.003 | 19.6 | 0 | 0.00 |
| Session 20181128 S23.set | 128 | 1000 | 197 | 0.002 | 12.9 | 0 | 0.00 |
| Session 20181129 S24.set | 128 | 1000 | 198 | 0.001 | 12.3 | 0 | 0.00 |
| Session 20181129 S25.set | 128 | 1000 | 169 | 0.003 | 17.7 | 0 | 0.00 |
| Session 20181211 S29.set | 128 | 1000 | 208 | 0.001 | 17.7 | 0 | 0.00 |
| Session 20181211 S30.set | 128 | 1000 | 192 | 0.002 | 14.8 | 0 | 0.00 |
| Session 20181211 S31.set | 128 | 1000 | 154 | 0.006 | 27.4 | 0 | 0.10 |

## 3. PIPELINE VALIDATION

To validate APICE, we compared it with a standard preprocessing pipeline (STD). In the Standard pipeline, bad channels were identified on segmented data using absolute thresholds. In APICE, the artifact rejection and correction were performed on continuous data. To identify the best level for the relative thresholds used during artifact detection, we run APICE with three different values: 2, 3, and 4 interquartile ranges (see section 2.2). With the intermediate relative threshold, 3, we also applied some variations to the APICE pipeline. In APICEa, we only applied the artifacts’ detection in the continuous data and avoided interpolating localized artifacts to evaluate the effect of the interpolation. Additionally, we applied W-ICA and DSS filters to evaluate if these cleaning algorithms improve data quality (Figure 1).

To have a more robust validation, we analyzed two very distinct datasets. Significant changes occurred in the EEG features during development (Eisermann et al., 2013; Marshall et al., 2002; Nelson & Monk, 2001), and the level of contamination by motion artifacts can vary considerably according to infants’ age and the type of task. Therefore, we decided to use two datasets differing in the age of the infants, in the modality of the stimulation, and in the type of task. The first dataset corresponds to an auditory experiment in neonates. Neonates were tested while sleeping, and the task consisted of passive listening, meaning that the data were minimally contaminated by motion artifacts. The second experiment corresponded to a visual task in which 5-month-old infants looked to a sequence of images on the screen. The infants were awake and actively engaged in the task, resulting in strong motion artifacts in the data.

### 3.1. Datasets

#### 3.1.1. Dataset 1: Neonates dataset

The neonate’ dataset corresponds to an auditory experiment. During each trial, infants heard 4 or 5 syllables lasting 250 ms presented every 600 ms. 216 trials were presented to each infant. Scalp electrophysiological activity was recorded using a 128-electrode net (Electrical Geodesics, Inc.) referred to the vertex with a sampling frequency of 250 Hz. Neonates were tested in a soundproof booth while sleeping or during quiet rest. Participants were 24 (11 males) healthy-full-term neonates, with normal pregnancy and birth (gestational age > 38 weeks of gestation, Apgar score ≥ 7 in the first minute and ≥ 8 in the fifth minute, and cranial perimeter ≥ 33.0cm). All participants were tested at the Port Royal Maternity (AP-HP), in Paris, France. Parents provided informed consent.

#### 3.1.2. Dataset 2: 5-month-old dataset

The 5-month-old infants’ dataset was a study investigating their capacity to associate two sets of images. During each trial, an attractor appeared on the center of the screen for 0.6 s, followed by a first image lasting 1 s, a second image lasting 1.2 s, and the attractor again during other 1.0-1.2 s. The experiment lasted until the infants were fussy (80-140 trials per participant). Scalp electrophysiological activity was recorded using a 128-electrode net (Electrical Geodesics, Inc.) referred to the vertex with a sampling frequency of 500 Hz. Infants were tested in a soundproof shielded booth while sitting in their parents’ lap. Participants were 26 (12 males), 22.98-weeks-old infants (SD 1.41, min 20.86, max 27). All participants were tested at NeuroSpin, in Gif/Yvette, France. Parents provided informed consent

### 3.2. Preprocessing

We included different preprocessing approaches. The final steps were the same for all methods. After bad epochs were removed and data were average referenced, data were baseline corrected using the average over [-100, 100] ms.

#### 3.2.1. Standard pipeline

In the standard pipeline, the data were filtered (low-pass filter at 40 Hz and high-pass filter at 0.2 Hz) and epoched. Afterward, we defined bad channels per epoch based on three criteria. First, we discarded channels with a low (below 0.4) top 5 % correlation with the other channels. Then, we rejected channels with an amplitude bigger than 500 mV on non-average reference data. Finally, we rejected channels with a fast running average bigger than 250 mV or a difference between the fast and slow running average bigger than 150 mV on average reference data. If less than 30 % of the channels were rejected, they were interpolated using spherical splines. If more than 30 % of the channels contained artifacts, the epoch was rejected. The retained trials were average referenced and baseline corrected using the average over [-100, 100]. All trials were averaged in each infant and then across infants to create a grand average ERP.

#### 3.2.2. APICE pipeline

The APICE pipeline consisted of the steps described in each of the corresponding sections above. Data were filtered (low-pass filter at 40 Hz and high-pass filter at 0.1 Hz), and artifacts were detected on the continuous data (see section 2.2). Afterward, bad samples and channels were defined (see section 2.3). Bad samples were identified as those with more than 30 % of the good channels rejected and lasting at least 100 ms. Bad channels were those presenting artifacts during more than 30 % of the good samples. Artifacts were corrected using target PCA on segments shorter than 100 ms and using spatial spherical spline to interpolate bad channels (see section 2.4). Finally, artifacts were detected again, and bad samples and channels were re-defined.

We used three different relative thresholds. As described in section 2.2, the relative thresholds are fixed based on a certain number of interquartile ranges from the first and third quartiles. We fixed this value to 2 (APICE (2)), 3 (APICE (3)), and 4 (APICE (4)) in different runs of the preprocessing to evaluate how it affects the rejection percentage. A lower value (2) detects more artifacts but discards more data. A higher value (4) misses some artifacts but keeps more data.

To obtain ERPs, the continuous preprocessed data was further high-pass filtered at 0.2 and epoched. Then, bad samples and channels were re-defined on the epoched data based on the data already rejected. A sample was defined as bad as explained on continuous data. A channel in a given epoch was defined as bad if it presented any artifact lasting more than 100 ms. Notice that some channels may present artifacts event during periods not defined as bad times, because we re-detected artifacts after the correction of transient artifacts (see Figure 2D). If less than 30 % of the channels were bad, they were interpolated using spherical splines. Epochs were rejected based on the amount of bad data. Either when more than 30 % of the channels were bad channels or when the epoch contained any bad time. Finally, the rejected epochs were removed, data was average referenced, and the average over the period [-100, 100] ms was used as the baseline. All trials were averaged in each infant and then across infants to create a grand average ERP.

#### 3.2.3. APICE+W-ICA pipeline

The APICE +W-ICA pipeline was primarily the same as the APICE(3) pipeline using a relative threshold equal to 3 for the artifacts rejection steps. The only difference was that target-PCA was not applied during artifact correction because ICA should isolate the artifacts corrected by this algorithm. After artifacts identification and correction in the continuous data, ICA was applied as described in section 2.5. The steps to obtain the ERP are the same as in the APICE pipeline.

#### 3.2.4. APICE+DSS pipeline

The APICE +DSS pipeline is the same as the APICE(3) pipeline with a relative threshold equal to 3 for the artifacts rejection steps. The only difference is that after bad epochs were removed and the data were average-referenced, the DSS filter was applied to the remaining trials. In the first PCA, we retained 50 components, and in the second PCA 15. Finally, data were baseline corrected as in the other pipelines.

### 3.3. Pipelines evaluation

We report different metrics to compare pipelines performance. The metrics include: (1) the standardized measurement error (SME) to quantify the noise (Luck et al., 2020) applied to the auditory ERP for Dataset 1 and to the visual ERP for Dataset 2; (2) the proportion of retained epochs; and (3) the amount of interpolated data in the retained epochs.

The SME was computed for the average response over a certain region of interest and time window corresponding to the auditory or visual ERPs using bootstrapping. We randomly sampled with replacement N responses for each subject, where N is the number of trials retained. Then, we computed the mean in time and space. We repeated the process 1000 times, and the standard deviation of the measure across all iterations corresponded to the SME for each subject ERP (Luck et al., 2020). Higher SMEs denote nosier data and smaller SMEs cleaner data. The SME for Dataset 1 was computed over central electrodes in the time window 250-350 ms (Figure 4), which corresponds to the auditory response (Dehaene-Lambertz & Pena, 2001). The SME for Dataset 2 was computed over occipital electrodes in the time window 550-650 ms (Figure 4), which corresponds to the P400 visual ERP (de Haan & Nelson, 1999).

**Figure 4.**
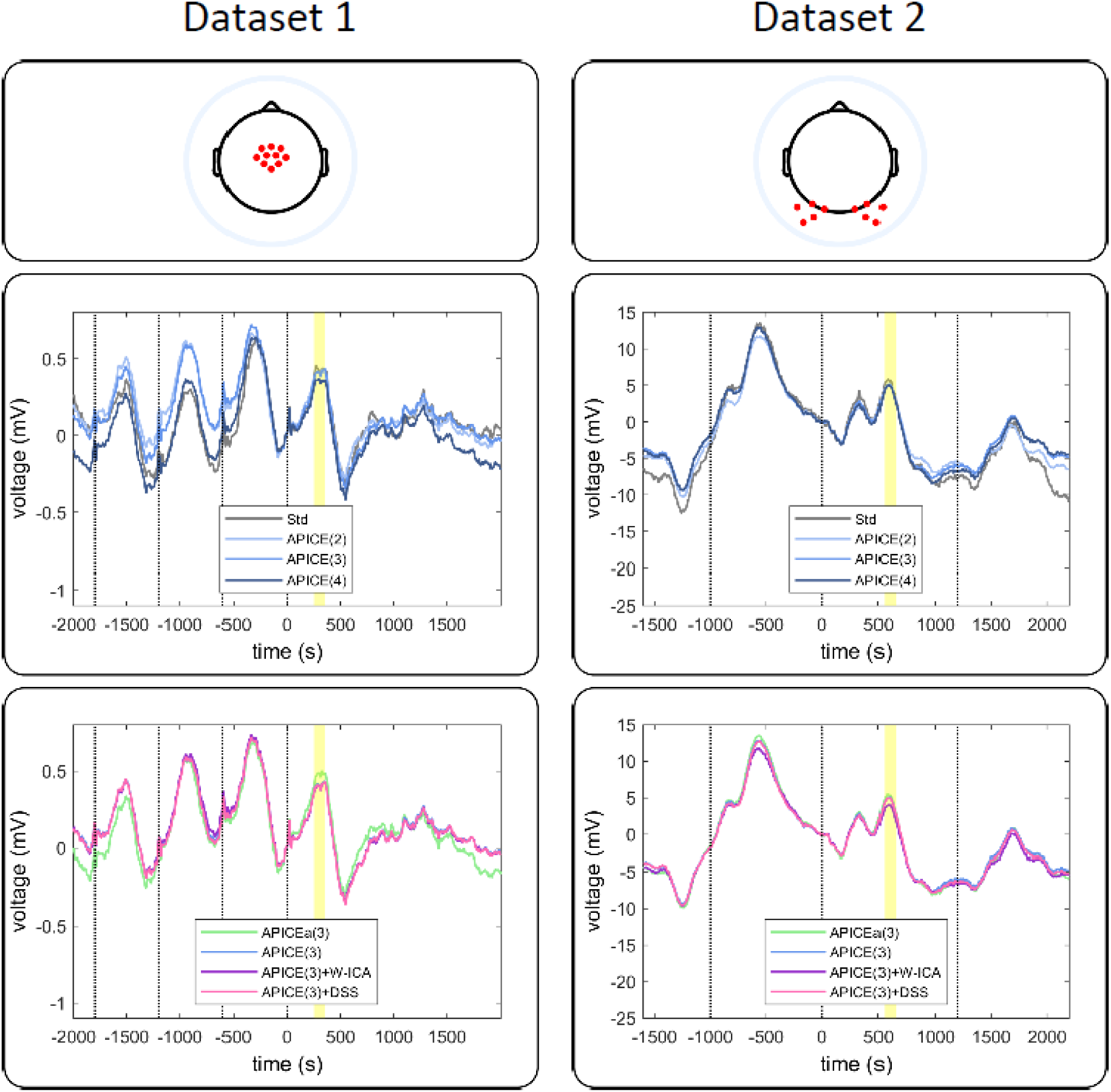
Grand average ERPs for datasets 1 and 2. The electrodes considered are indicated in red on the top panels. The shaded area shows the time windows when the SME was computed. The middle panel shows the ERP after preprocessing with the Standard pipeline and APICE pipeline with the three threshold levels for artifacts detection. The bottom panels shows the ERPs after preprocessing with the APICE(3) pipeline (relative threshold=3) and the pipelines incorporating also ICA and DSS.

### 3.4. Results

The ERPs obtained with the different methods on the two datasets are shown in Figure 4. As can be observed, the shapes of the ERPs do not seem to be affected by the applied preprocessing pipeline.

We first compared SME, percentage of retained epochs, and percentage of interpolated data obtained after applying the Standard and the APICE pipelines with the three artifact-rejection levels. For Dataset 1, the SME differed across pipelines (F(3,69) = 34.49; p = 9.83×10^-14^). Pairwise comparisons, Bonferroni corrected, show that it was larger (i.e. noisier data) for the Standard pipeline than for the APICE(2) (p = 4.0×10^-6^), APICE(3) (p = 6.1×10^-5^), and APICE(4) (p = 4.9×10^- 4^). The SME was lower in APICE(2) than in APICE(4) (p = 1.7×10^-5^) and APICE(3) (p = 8.6×10^- 5^), and in APICE(3) than in APICE(4) (p = 0.00249) (Figure 5A).The amount of retained data significantly differed across pipelines (*F(3,69) = 69.16*; *p < 2×10^-16^*). Pairwise comparisons, Bonferroni corrected, show that more epoch were retained in the Standard pipeline that in APICE(2) (p = 2.5×10^-8^), APICE(3) (p = 1.0×10^-6^), and APICE(4) (p = 1.5×10^-4^). As expected, more trials were retained in APICE(4) than APICE(2) (p = 9.5×10^-8^) and APICE(3) (p = 7.6×10^- 7^), and more trials were retained in APICE(3) than APICE(2) (p = 1.6×10^-7^) (Figure 5B). The amount of interpolated data differed as well (F(3,69) = 197.8, p < 2×10^-16^). Pairwise comparisons, Bonferroni corrected, show that less data was interpolated in the Standard pipeline than in the APICE(2) (p = 1.0×10^-12^), APICE(3) (p = 1.1×10^-10^), and APICE(4) (p = 2.6×10^-10^). Interpolation was more frequent in APICE(2) than APICE(3) (p = 3.2×10^-12^) and APICE(4) (p = 3.2×10^-12^), and in APICE(3) than in APICE(4) (p = 1.4×10^-9^) (Figure 5C).

**Figure 5.**
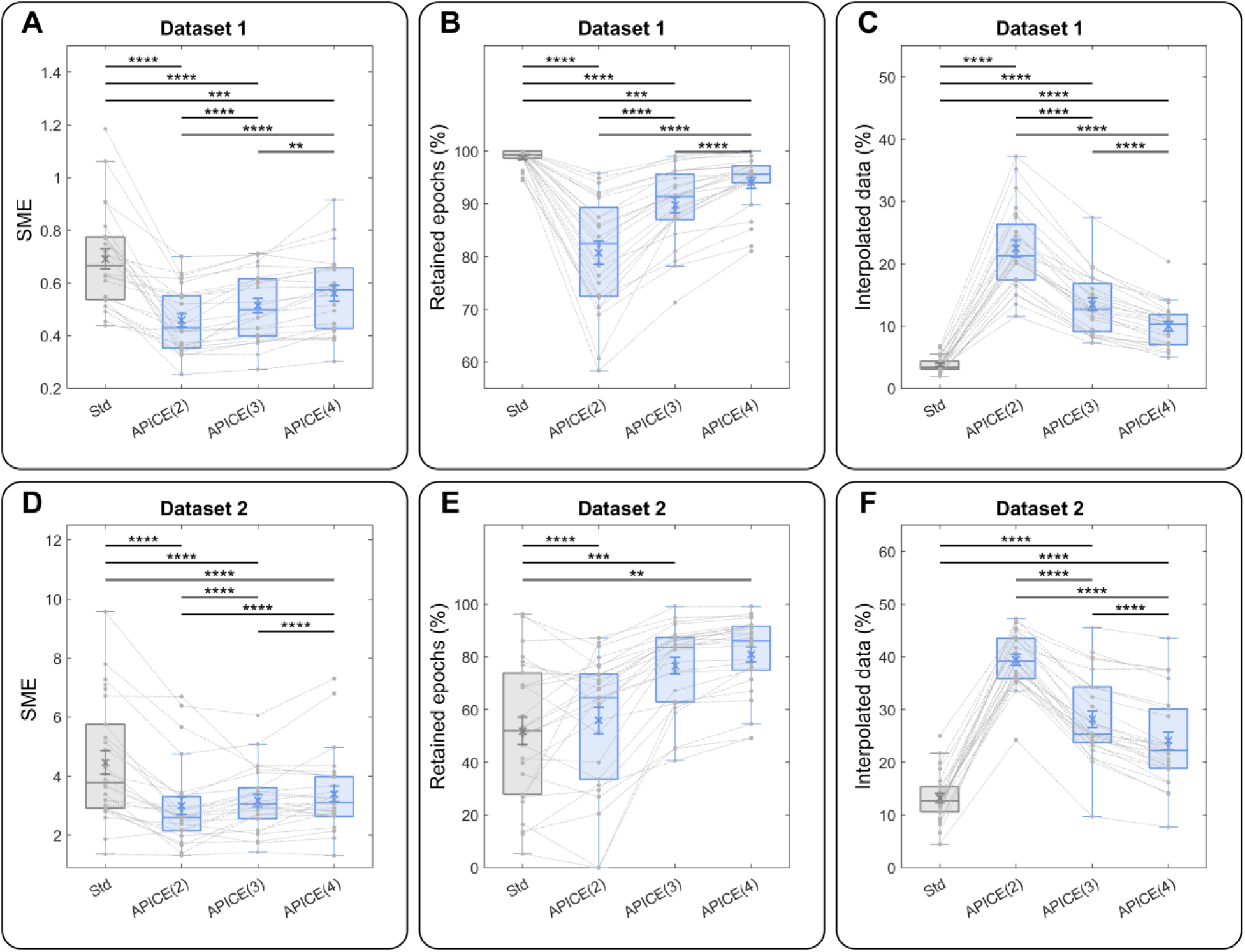
Comparison between the standard and APICE pipelines with three thresholds levels for artifact detection. **(A)** SME for Dataset 1. Higher SME indicates noisier data. **(B)** Percentage of retained epochs for Dataset 1. **(C)** Interpolated data for Dataset 1. **(D)** SME for Dataset 2. **(E)** Percentage of retained epochs for Dataset 2. **(F)** Interpolated data for Dataset 2.

For Dataset 2, the SME differed across pipeline (F(3,73) = 17.06, p = 1.7×10^-8^). Pairwise comparisons, Bonferroni corrected, show that the SME was larger for the Standard than for APICE(2) (p = 2.2×10^-5^), APICE(3) (p = 8.6×10^-4^), and APICE(4) (p = 0.00224). There was no significant difference between APICE(2), APICE(3), and APICE(4) (p > 0.1) (Figure 5D). The amount of retained epochs significantly differed across pipelines (F(3,75) = 39.43; p = 2.1×10^-15^) (Figure 5E). Pairwise comparisons, Bonferroni corrected, show no difference in the percentage of retained epochs between the Standard pipeline and APICE(2) (p > 0.1), but more epochs were retained in APICE(3) (p = 3.0×10^-6^) and APICE(4) (p = 7.2×10^-7^) than in the standard pipeline. As expected the proportion of epochs retained was larger in APICE(4) than APICE(2) (p = 1.2×10^-6^) and APICE(3) (p = 6.2×10^-5^), and in APICE(3) than APICE(2) (p = 1.7×10^-6^). The amount of interpolated data also differed (F(3,73) = 277.2, p < 2×10^-16^) (Figure 5F). Pairwise comparisons, Bonferroni corrected, show that less data was interpolated in the Standard pipeline than in the APICE(2) (p < 2×10^-16^), APICE(3) (p = 3.3×10^-13^), and APICE(4) (p = 2.5×10^-9^). Less data was interpolated in APICE(4) than in APICE(3) (p = 5.1×10^-10^) and APICE(2) (p = 3.1×10^-12^), and in APICE(3) than APICE(2) (p = 1.3×10^-12^).

We then investigated whether the interpolation of localized artifacts brings any improvement to the APICE pipeline (Figure 6), and compared the three measures after APICE, our full pipeline, and APICEa, a reduced version of APICE in which only the artifacts detection of APICE was implemented. In Dataset 1, we observed no difference in the SME (t(23) = 0.057268; p = 0.9548); but more data were retained after APICE (t(23) = 6.038; p = 3.7×10^-6^; mean of the difference 3.84%), and more data were interpolated in epochs after APICE (t(23) = 3.2739; p = 0.0033; mean of the difference 1.22%). In dataset 2, with APICE we observed a slightly lower SME (t(25) = 2.4295; p = 0.02264; mean of the difference 0.23); more data retained with APICE (t(25) = 6.1227; p = 2.119×10^-6^; mean of the difference 4.84%); and no difference in the amount of interpolated data (t(25) = 1.0135; p = 0.3205).

**Figure 6.**
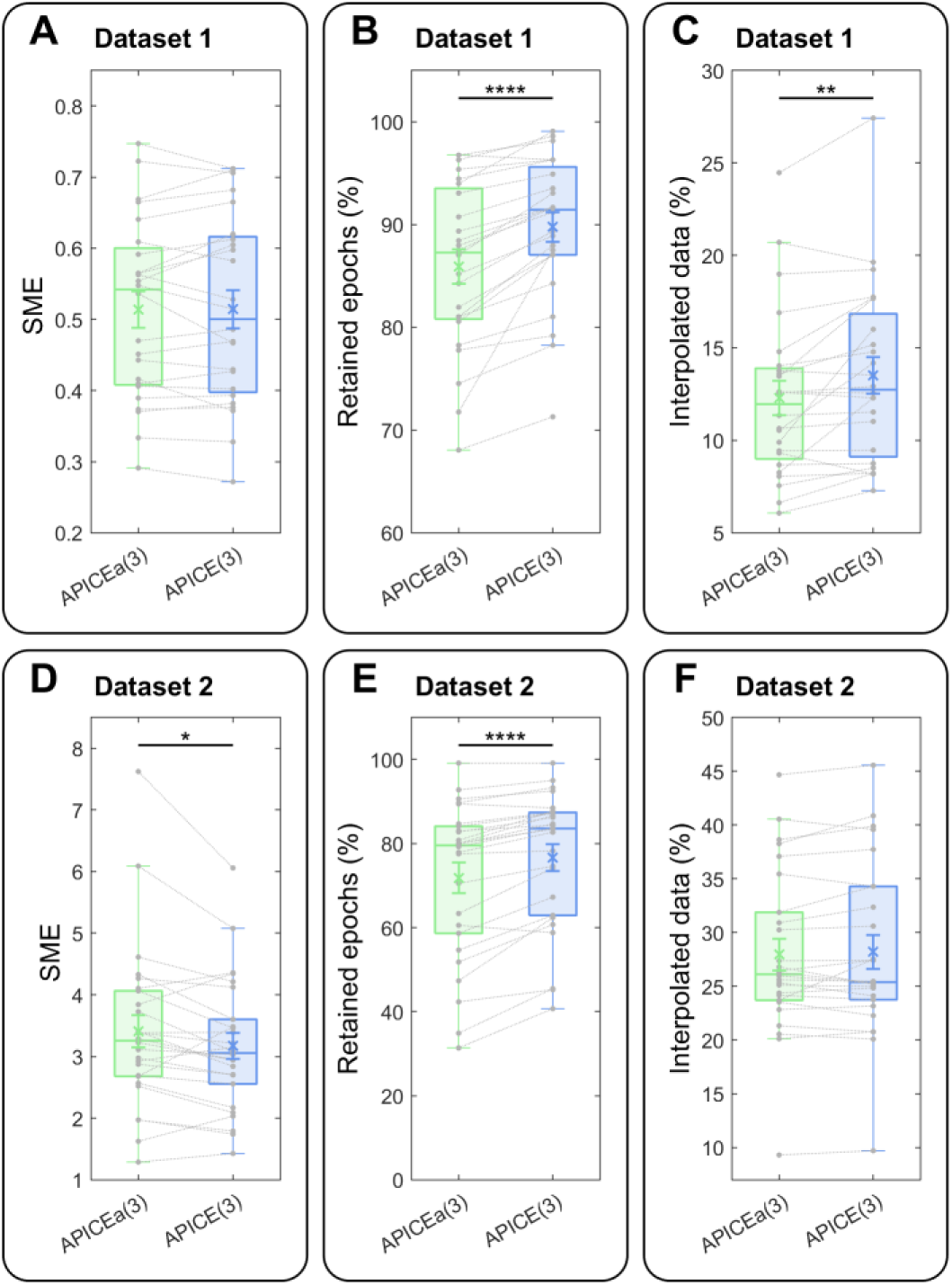
Comparison between our pipeline, APICE, and its reduced version, APICEa, in which artifacts were detected in the continuous data but without interpolation. **(A)** SME for dataset 1. **(B)** Retained epochs for dataset 1. **(C)** Interpolated data for dataset 1. **(D-E-F)** same than **A-B-C** for dataset 2.

Finally, we considered the two supplementary cleaning methods: ICA and DSS. The ICA step applied on Dataset 1 removed on average 11.9% of the components (SD 8.37%, min 0%, max 30%) and 0.57% of the total variance (SD 0.46%, min 0%, max 2.09%). On Dataset 2, it removed on average 48.3% of the components (SD 17.18%, min 12%, max 75%) and 2.89% of the total variance (SD 1.71%, min 0.07%, max 7.30%). The DSS filter on Dataset 1 removed 13.18% of the total variance (SD 4.13%, min 6.51%, max 23.11%). On Dataset 2, it removed 33.46% of the total variance (SD 5.31%, min 19.48%, max 44.43%).

Neither of the data cleaning methods, ICA or DSS, made a critical change (Figure 7). In Dataset 1, ICA marginally improved the SME (t(23) = 2.0848; p = 0.04838; mean of the difference 0.00957), slightly decreased the amount of retained epochs (t(23) = 2.5841; p = 0.0166, mean of the difference 0.17%), and slightly decrease the percentage of interpolated data (t(23) = 3.5399; p = 0.001749; mean of the difference 0.095%). In Dataset 2, ICA slightly decrease the SME (t(25) = 2.3089; p = 0.0295; mean of the difference 0.1468), did not affect the amount of retained epochs (t(25) = 1.3872; p = 0.1776), and slightly decreased the amount of interpolated data (t(25) = 7.4111; p = 9.2×10^-8^; mean of the difference 0.47%). Meanwhile, the DSS slightly decreased the SME (t(23) = 3.3037; p = 0.003102; mean of the difference 0.0176) when applied to dataset 1,and did not affect the SME when applied to dataset 2 (t(25) = 0.96803; p = 0.3423).

**Figure 7.**
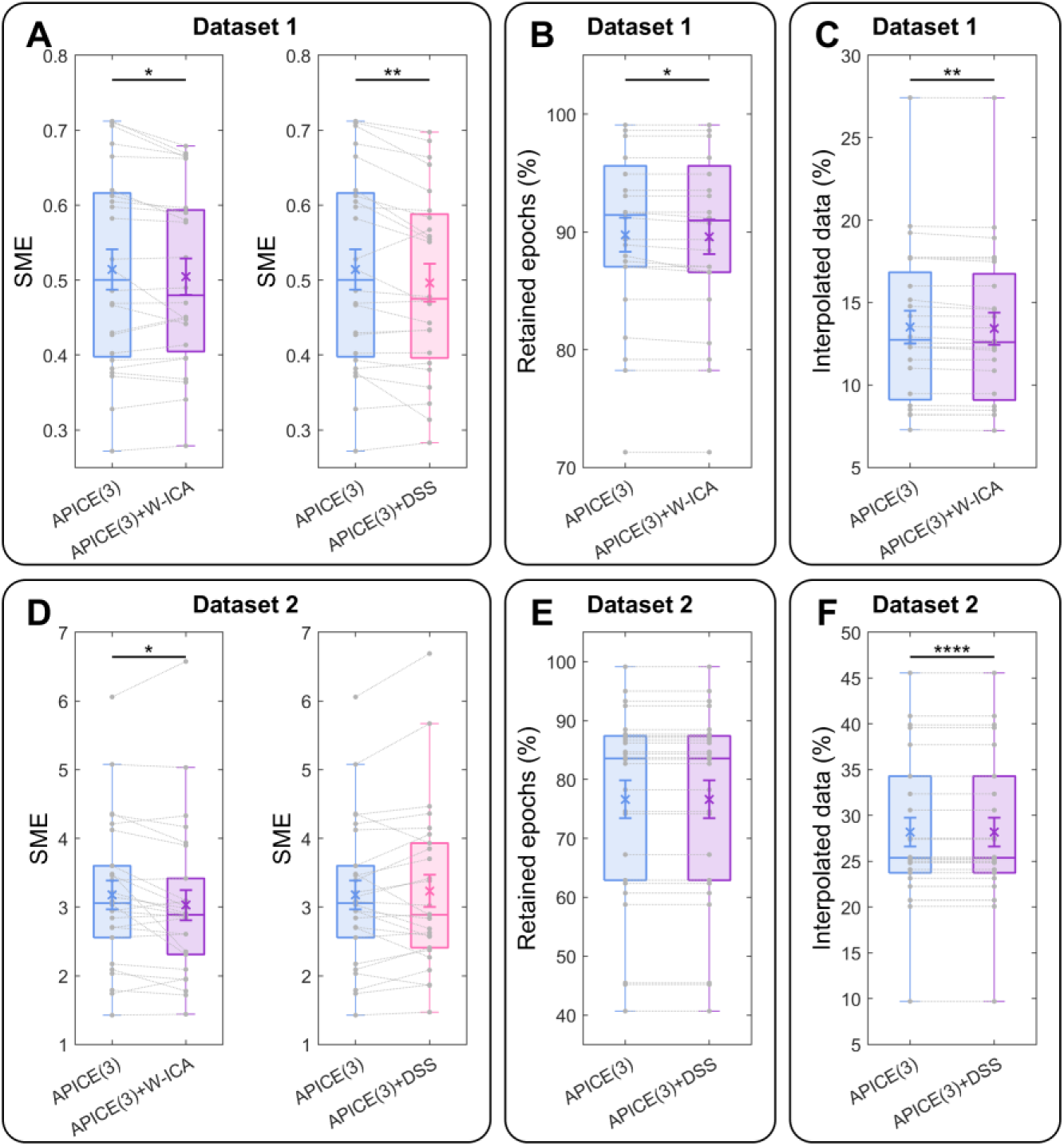
Comparison between APICE and APICE with the addition of ICA or DSS. **(A)** SME for dataset 1. **(B)** Retained epochs for dataset 1. **(C)** Interpolated data for dataset 1. **(D)** SME for dataset 2. **(E)** Retained epochs for dataset 2. **(F)** Interpolated data for dataset 2.

### 3.5. Discussion

Using two widely different infant datasets, we compared our pipeline with a standard preprocessing pipeline. In traditional pipelines, data is usually segmented after filtering, epochs containing artifacts are identified based on absolute thresholds (e.g., the absolute amplitude or a running average of the signal larger than a threshold), and bad channels are eventually interpolated. In the APICE pipeline, different algorithms based on relative thresholds are used to identify artifacts, and the detection and correction steps are performed on the continuous data. We evaluated the pipelines on the stability and reproducibility of the obtained ERPs through the SME procedure. This procedure consists of examining the distribution of the ERP values through multiple random draws of the trials used to compute the subject ERP. This value should be low if the averaging process correctly cancels the remaining and unwanted “noise”; that is to say if no unexpected large-amplitude not-evoked activity remains in the preprocessed data. We also examined the number of kept trials that can be important for complex paradigms in which pertinent conditions comprise only a few trials. We additionally report the proportion of interpolated data that corresponds to a loss of data, although this loss is relative given the high redundancy between spatial and temporal samples of EEG data.

Results show that the APICE pipeline outperforms the standard pipeline for both datasets. SMEs were smaller in both cases with all the tested relative thresholds levels. For Dataset 1 (an auditory experiment in sleeping neonates), which was only mildly contaminated by motion, the standard pipeline retained more trials than our pipeline. While more kept trials are, in principle, an advantage to recover the ERP through the average process, it is not anymore when it implies a diminished data quality. The neonates were tested asleep, which means that high amplitude artifacts exceeding the absolute thresholds of the standard pipeline were rare. However, implementing various algorithms based on different signal features makes our detection procedure more sensitive to outlier signals. In Dataset 2 (a visual experiment in awake 5-month-old infants), where the contamination by motion artifacts was substantial, APICE shows a smaller SME and a smaller amount of rejected data. Thus, relative thresholds determined by subject and electrode make the artifacts’ detection in APICE more accurate, resulting in improved data quality and better recovery of good quality data.

We also validated APICE on three levels for the relative threshold. Increasing the threshold from 3 to 4 hardly changes the percentage of rejected data, but the SME decreases. On the contrary, a reduction to 2 slightly reduces the SME but at the cost of a much higher rejection. Based on these results, we recommend using a threshold of 3 as default to keep enough epochs per subject. Nevertheless, the threshold can be adjusted depending on the analysis requirements –either more data but noisier or less data but cleaner. Remember that the ERP results from a compromise between an acceptable amplitude variation and keeping many trials to cancel non-evoked activity through averaging.

Besides performing artifacts detection on the continuous data, APICE also corrects localized artifacts. We tested if this step brings any improvement by comparing APICE with APICEa, a reduced version in which we removed the interpolation of transient artifacts. Results show that while in APICE, the SME is only marginally lowered relatively to APICEa, the amount of retained epochs is considerably higher. Thus, the interpolation of transient artifacts enables the recovery of otherwise lost trials without loss of data quality.

Altogether, our results demonstrate that using multiple algorithms with relative thresholds combined with the interpolation of transient artifacts applied on continuous data improves artifacts detection and data recovery relative to basic algorithms with fixed absolute thresholds. This approach also brings flexibility because the same preprocessed data can serve to perform analyses requiring different segmentation strategies. In addition, accurate detection of artifacts allows deciding how to handle them in subsequent processes. For example, we can exclude motion artifacts before performing ICA or use the amount of data with artifacts in an epoch as a criterion to reject it.

Finally, we also tested if ICA and DSS, two data cleaning methods extensively used in adult studies, improve data quality. For ICA, we implemented a procedure that automatically removes components identified as non-neural artifacts using iMARA (Marriot Haresign et al., 2021) from non-high passed filter continuous data (section 2.5). Results show a modest improvement (decrease in the SME) restricted to Dataset 2. Similarly, for DSS applied on the retained epochs, we observed a slight improvement in the data quality, particularly in Dataset 1. In brief, our results show no substantial improvement in data quality, neither for ICA nor for DSS, even though we tried the latest proposals (combination of ICA with wavelet-thresholding to achieve a better ICA decomposition (Rong-Yi & Zhong, 2005), as it was also carried out in HAPPE, another pipeline for developmental studies (Gabard-Durnam et al., 2018)). We also used iMARA (Marriot Haresign et al., 2021), a recent modification of MARA (Winkler et al., 2011), to adapt it to infant data. Despite our effort to optimize ICA, we did not see a clear benefice for infants’ ERP analysis. It is worth noticing that we only tested iMARA for the IC classification; thus, it is possible that other available methods, like adjusted-ADJUST (Leach et al., 2020), may provide better performance.

One possible explanation is that these methods are not optimal for infant data due to the intrinsic properties of the signal. For example, the high inter-trial variability observed in infants (Naik et al., 2021) might complicate the separation of variance related to evoked and non-evoked activity, as the method is based on the presence of highly reproducible activity across trials. The developmental changes and variability in infant EEG neural responses and physiological artifacts might also result in a poor decomposition during ICA for infant data. Thus, further optimization of the algorithms is needed to achieve better performance. While iMARA performs better than the original MARA algorithm on the classification of non-neural IC (Marriot Haresign et al., 2021), its modifications are in the computation of the features used for classification. However, the classification is still based on the same adult learning set. Given the negligible improvement in the noise level and the fact that ICA is very time-consuming, we do not consider its application justified in the current state of the art. Concerning DSS, we do not recommend its application unless numerous trials are available.

## 4. CONCLUSION

EEG is a widely used technique in developmental studies. Nevertheless, there is no standardization of infant data preprocessing, partly because of a smaller community than for adult EEG/MEG and partly because of the many challenges that preprocessing infant data entails. Infants’ recordings are not only shorter but also more heavily contaminated by motion artifacts. Moreover, the types of artifacts and the features of the EEG signal change during development (Eisermann et al., 2013; Kushnerenko et al., 2002; Marshall et al., 2002; Nelson & Monk, 2001), making even harder the design of methods applicable across age groups. Consequently, most of the approaches optimized for adult data are ineffective on infant data.

When preprocessing infant data, it is fundamental to remove segments contaminated by motion artifacts before applying ICA or any other cleaning step to guarantee good performance. APICE successfully identifies artifacts across different ages and experimental conditions by using multiple algorithms and adaptive thresholds. Nevertheless, crucial challenges remain. Many physiological artifacts have amplitude and spectral properties that make them indistinguishable from the neural signal (e.g., eye movements, muscle artifacts) by algorithms looking at the local properties of the signal. For example, in APICE, we can identify the heartbeat using an algorithm detecting fast changes in the signal, but APICE does not explicitly search for artifacts like blinks or eye movements. Moreover, rejecting any segment contaminated by physiological artifacts would cause an enormous loss of data. In APICE, we partially deal with this problem by applying a target PCA to transient artifacts. In this way, we can partially remove heartbeats and jumps in the signal. However, more complete and efficient removal of all physiological artifacts would require blind source separation methods (Islam et al., 2016; Jiang et al., 2019), and the algorithms proposed in adults are, for the moment, not satisfactory. A better description of infants’ data features is necessary to design more suitable methods.

We created APICE to be fully automated and flexible. Automation guarantees replicability and scalability for growing data sets (without increasing the human workload). To ensure flexibility and better data recovery, APICE performs all the artifact detection steps in the continuous data. Consequently, the same preprocessed data is ready for different types of analysis. Furthermore, APICE is modular, allowing it to be easily modified to meet specific needs and incorporate new preprocessing steps. APICE also includes functions for renaming events, correcting their timings, organizing trials by condition, and computing the average ERP. APICE is freely available at https://github.com/neurokidslab/eeg_preprocessing, with example scripts illustrating its application. APICE is currently limited to MATLAB, but we want to make it available in an open-source programming language in the future, especially to integrate it with MNE python.

## Authors Contribution

A.F. and G.D.L. conceived the pipeline. A.F. wrote the software. G.G. and L.B. contributed to test the software. A.F. wrote the manuscript. G.G., L.B., and G.D.L. revised the manuscript and the pipeline validation.

## Acknowledgments

This research has received funding from the European Research Council (ERC) under the European Union’s Horizon 2020 research and innovation program (grant agreement No. 695710). We thank the families that participated in the studies. We would also like to thank Marie Palu and Chanel Valera for their help in recruiting and testing the infants and the entire Neurokids group at Neurospin for their feedbacks.

## APPENDIX A Artifact detection algorithms

The rejection matrix is stored in EEG.artifacts.BCT.

The input of each artifact algorithm is the EEGLAB structure and some parameters. The output of each artifact detection algorithm is the EEGLAB structure and a logical matrix containing the data rejected by that algorithm. The algorithms update the rejection matrix in the EEGLAB structure, such as it contains the old and new rejected data. The data already rejected is not used for the estimation of relative thresholds.

The following algorithms are particularly sensitive to the detection of non-functional channels:

- ***eega_tRejCorrCh:*** This algorithm relies on the high correlation existing between adjacent channels, especially in high-density systems. The correlation between all channels is computed in sliding time windows (4 s length, 2 s step by default), and for each channel and time window, the average over the stronger correlations (top 5% by default) is the measure used for defining bad data. The algorithm rejects channels per time window with a top correlation lower than a threshold. Because the method’s measure is independent of the signal amplitude and its distribution is not normal, we recommend using an absolute threshold (0.4 by default). However, the function supports the estimation of a unique relative threshold for all electrodes.
- ***eega_tRejPwr:*** This algorithm identifies segments of bad data based on the power at different frequency bands. It applies FFT in sliding time windows (4 s length, 2 s step by default), and computes the average power in each frequency band, electrode, and time window, and expresses it in decibels relative to the median value across all time windows and electrodes. By default, the algorithm z-scores the data per channel, uses relative thresholds, and rejects data when the power in the frequency band [1, 10] Hz is below the threshold or the power in the frequency band [20, 40] Hz is above the threshold. Notice that for this algorithm, the same threshold is used for all electrodes.

The following algorithms are more specifically sensitive to motion artifacts, but they will also partially identify non-working channels and other types of artifacts.

- ***eega_tRejAmp:*** This algorithm rejects samples with an amplitude below or above a threshold. By default, it sets the thresholds relatively and per electrode and applies a 50 ms mask (it removes the 50ms before and after any segment rejected by the algorithm).
- ***eega_tRejTimeVar:*** This algorithm computes the variance of the signal in a sliding time window (0.5 s length, 0.1 s step by default) and rejects samples with a variance above or below a threshold. By default, it sets the thresholds relatively and per electrode.
- ***eega_tRejRunningAvg:*** This algorithm computes for each sample two weighted running averages and rejects samples with a too high fast running average or with a too high difference between the fast and the slow running average. The fast-running average is computed as *AvgF_j_* = 0.800 × *AvgF_j_*_−1_ + 0.200 × *X_j_* where *X_j_* is the data at sample *j*, and the slow running average as *AvgS_j_* = 0.975 × *AvgF_j_*_−1_ + 0.025 × *X_j_*. By default, it sets the thresholds relatively and by electrode and applies a 50 ms mask to the bad segments.
- ***eega_tRejFastChange:*** This algorithm rejects data when the maximum change in a given time window (20 ms by default) is larger than a threshold. It specifically detects jumps in the signal and can identify some non-neural activity like heartbeats. By default, it sets the thresholds relatively and by electrode.
- ***eega_tRejAmpElecVar:*** This algorithm rejects data based on the variance of the signal across electrodes. If the amplitude for a given electrode is too far away from the median of all electrodes, it is rejected. Notice that for this algorithm, the threshold is by definition relative and defined for all the electrodes. By default, it applies a 50 ms mask to the bad segments.

The following algorithms can be used to reject/re-include data according to the rejection matrix *EEG.artifacts.BCT*.

- ***eega_tIncShortBad:*** This function includes segments of rejected data that are shorter than a lower threshold (20 ms by default).
- ***eega_tRejShortGood:*** This function rejects segments of good data between segments of bad data if they are shorter than a lower threshold (2 s by default).
- ***eega_tMask:*** This function applies a mask to the bad segments (50 ms by default).
- ***eega_tRejChPercSmpl:*** This function rejects all the data for channel and epoch if more than an upper limit of samples were rejected (50% by default).
- ***eega_tRejSmplPercCh:*** This function rejects all the data at a particular time-point if more than an upper limit of channels were rejected (30% by default).

We also provide a function to run a combination of multiple artifact detection algorithms.

- ***eega_tArtifacts:*** This function runs a combination of artifact detection algorithms to ensure proper artifact detection. The function performs a defined number of loops of rejection. Within each loop, it applies the specified algorithms on the data remaining from the previous loops. After each loop, it updates the rejection matrix. Both the algorithms to run and its parameters are provided to the function as a structure. For some algorithms to work correctly, it is important that the data is filtered—especially low-pass filtered to remove line noise. It can be specified that the data needs to be low or high pass filter before performing the artifact rejection (notice that the function never modifies the output data, the filtered data are only used for the computation). The function can either reset or keep (default) the initial rejection matrix.

## APPENDIX B Description of the function to define BT and BC

We created a single function to define bad times (BT), bad channels per epoch (BC), and bad channels during the whole recording (BCall) on both continuous and epoched data. Notice that the definition of bad times and channels based on the rejection matrix are linked. For example, if we define BT as samples with more than 15% of rejected electrodes and that for one subject, 18% of the channels are non-functional, we will define all times as bad and discard the entire recording. To avoid this kind of problem, we created a function that can iteratively approximate the final thresholds used for defining BT, BC, and BCall.

- ***eega_tDefBTBC:*** This function defines *EEG.artifacts.BT* as a logical matrix with size 1 x samples x epochs signaling the bad samples, *EEG.artifacts.BC* as a logical matrix with size channels x 1 x epochs specifying the bad channels during each epoch, and *EEG.artifacts.BCall* as a logical vector with a length equal to the number of channels, indicating non-functional channels for the entire recording. The thresholds used to define BT, BC, and Bcall have to be provided as input. If more than one threshold is provided for each definition, they will be applied as follows. First, the function defines bad times as those where the proportion of rejected channels (excluding bad-channel in BC) at each sample is above the first threshold. Second, it defines bad channels during the whole recording as those for which the proportion of total rejected samples (excluding bad times in BT) is above the first threshold. Third, it defines bad channels per epoch in an analog way, but computing the rejected samples per epoch and using the corresponding first threshold. It repeats the process by looping over all the thresholds provided. This iterative approximation to the thresholds allows a better definition of BT, BC, and Bcall. The function also allows applying a mask to bad-times (non applied by default), ignoring bad times shorter than a lower limit (0 by default), and marking as bad-times short good-times between two bad-times (0 by default).

## APPENDIX C Description of the artifact correction algorithms

We provide a function to perform the target PCA correction and two separate functions to perform the spatial interpolation of channels. One function interpolates channels non-functional during the entire recording (or an entire epoch when applied on epoched data). The second function interpolates segments of rejected data. All functions mark corrected data as good in the rejection matrix and indicate which data has been interpolated in another logical matrix, *EEG.artifacts.CCT* (true indicates corrected data).

- ***eega_tTargetPCAxElEEG*:** This functions corrects transient artifacts using target PCA. It concatenates the data segment to correct, applies PCA to this subset of data, and removes the first *n* components. Finally, the interpolated segments are spliced back into the data and aligned to it to avoid discontinuities. By default, the segments are concatenated relatively to the immediate previous sample. Note that this alignment introduces drifts in the signal; thus, after artifact correction using this method, the data should be high-pass filtered again. By default, the correction is restricted to data segments indicated as good data in *EEG.artifacts.BT*, shorter than 100 ms, and to the channels indicated as good in *EEG.artifacts.BC*.
- ***eega_tInterpSpatialSegmentEEG*:** This function uses spherical spline to spatially interpolate channels not working during limited periods. The interpolated segments are spliced back into the data and aligned to it to avoid discontinuities. By default, the segments are concatenated relatively to the immediate previous sample. As before, this alignment introduces drifts in the signal, requiring a subsequent high-pass filtering. The signal is reconstructed only during segments indicated as good data in *EEG.artifacts.BT* and when the proportion of channels rejected in the rejection matrix is lower than a threshold. By default, segments shorter than 100 ms are not interpolated, and a 1 s mask is applied to all segments before interpolation.
- ***eega_tInterpSpatialEEG:*** This function uses a spherical spline to interpolate channels identified in *EEG.artifacts.BC* as non-functional during an entire epoch (or the entire recording when applied to continuous data).

## APPENDIX D Description of the function for ICA and DSS

We created a function that performs the ICA combined with wavelet-thresholding ICA and afterward uses the iMARA algorithm to identify components associated with non-neural artifacts automatically. For short recordings, the function can run the analysis on subsets of electrodes or use PCA to reduce the dimensionality.

- ***eega_pcawtica:*** This function applies ICA to the removal of non-neural sources. The function operates as follows. *(1)* It makes a copy of the data, which is high-pass filtered (at 2 Hz by default) and low-pass filter (at 40 Hz by default). *(2)* It removes bad samples and bad channels. *(3)* If channel subsets are provided, it restricts the analysis to them. *(4)* It performs a first PCA (optional) + ICA. *(5)* It uses wavelet-thresholding to estimate transient artifacts and subtracts them from the data. *(6)* It performs a second PCA (optional) + ICA on the artifacts-free data. *(7)* It identifies the components to remove using iMARA. *(8)* It estimates the artifacts using those components. *(9)* It removes the estimated artifacts from the original data.

We provide a function, dss_denoise_EEG, which performs a DSS on an EEGLAB structure containing epoched data.

- ***dss_denoise_EEG:*** This function performs a DDS. It applies a first PCA, discards the last components, and normalizes the retained components. Then, it computes the average ERP, applies a second PCA to the average. The filter is the conjunction of the two PCA decompositions. Afterward, it projects the data into the filter and keeps only the first components. Finally, it projects the data back to the sensor space. This function enables designing a single filter and applying it to all the epochs or designing different filters for each condition (in this case, experimental factors have to be previously defined using *eega_definefactors*). The number of components to retain in the first and second PCA are provided as inputs to the function.

## APPENDIX E Other functions

Usually, a delay exists between the time the stimulation computer gives the order of presenting a stimulus and its actual presentation. Therefore, the latency of the events is usually corrected by the latency of DINs. We provide a function that performs this latency correction.

- ***eega_latencyevent:*** This function corrects the latency of certain types of events (indicated as input) using the latency information of the first DIN event (its name also has to be indicated as input).

For segmenting the data, we provide a function that, besides epoching the data, also epochs the logical matrices defining artifacts.

- ***eega_epoch:*** This function epochs the data and the logical matrices in the *EEG.artifacts* field.

We also provide two functions for identifying bad epochs, either based on the amount of rejected data or on the distance to the average evoked response.

- ***eega_tDefBEbaddata:*** This function identifies bad epochs based on the amount of rejected data and saves it as a logical vector in *EEG.artifacts.BE*. The function defines epoch *k* as bad based on *(1)* the proportion of bad data in *EEG.artifacts.BCT(:,:,k)*, *(2)* of bad samples in *EEG.artifacts.BT(1,:,k)*, *(3)* of bad channels in *EEG.artifacts.BC(:,1,k)*, and *(4)* of corrected data in *EEG.artifacts.CCT(:,:,k)*.
- ***eega_tDefBEdist*:** This function identifies bad epochs based on the Euclidean distance of the average referenced response during epoch *k* to the average response across all epochs. It calculates the Euclidean distance to the average at each sample for all epochs. Then it establishes a threshold based on a certain number of interquartile ranges from the third quartile. If for epoch k, the distance to the average response is bigger than the threshold during a certain proportion of the epoch, the function rejects the epoch.

After data has been segmented and the epochs with artifacts rejected, additional steps are usually performed to obtain the ERP responses. These typically include average referencing, and eventually, data normalization and baseline correction. While normalizing the data is not necessary, this process can improve statistical power. We provide a function for data normalization flexible to normalized base on different dimensions.

- ***eega_normalization:*** This function z-scores the data. The function can apply the normalization over single or all electrodes and single or all epochs. It is also possible to define specific values for the mean or the standard deviation. For example, setting the mean to 0 implies only dividing by the standard deviation.

In classical ERPs analysis, the responses across trials belonging to the same experimental condition are averaged to obtain an ERP per condition and subject. While the selection of trials usually needs to be adapted to specific requirements of each experiment, we provide two functions that can facilitate this process. The first defines experimental factors across trials, and the second averages across any of them.

- ***eega_definefactors*:** This function creates a field, *F*, in the EEG structure that specifies different experimental factors. *F* is a cell array containing all the possible experimental factors. The values for each experimental factor are determined based on the information present within each epoch in *EEG.epoch(i).(field of interest)*.
- ***eega_avgdatabyfactors:*** This function averages across one or multiple of the factors previously defined.

To obtain a summary of the preprocessing output for a group of subjects, we created a function that prints a table with the number of channels, samples, and epochs, the number of rejected and interpolated points, and the number of bad times, bad channels, and bad epochs. The table is created for some critical points during the pipeline: before epoching the data, before the rejection of bad epochs, and on the final stage

- ***eega_printsummary:*** This function takes all the files in a given folder adhering to a specific name and creates a summary report for them.

When preprocessing a dataset, a series of processes are sequentially applied to many files. We have created a function that sequentially applies a series of specified functions to a set of files. The function operates on all the specified files in an input folder and saves the final files in an output folder.

- ***eega_RunAll:*** This function considers all the files adhering to a provided file name in a specified folder and saves the resulting EEGLAB structure in a specified output folder.

The functions to apply and their parameters are provided to the function as a string, corresponding to the function name, followed by a cell array containing its inputs (excluding EEGLAB structure). It can run the functions for all the files found in the input folder or limit the output files that do not exist. It can create the output name by concatenating a string provided as input with the input file name.

